# Benchmarking solutions to the T-cell receptor epitope prediction problem: IMMREP22 workshop report

**DOI:** 10.1101/2022.10.27.514020

**Authors:** Pieter Meysman, Justin Barton, Barbara Bravi, Liel Cohen-Lavi, Vadim Karnaukhov, Elias Lilleskov, Alessandro Montemurro, Morten Nielsen, Thierry Mora, Paul Pereira, Anna Postovskaya, María Rodríguez Martínez, Jorge Fernandez-de-Cossio-Diaz, Alexandra Vujkovic, Aleksandra M. Walczak, Anna Weber, Rose Yin, Anne Eugster, Virag Sharma

## Abstract

Many different solutions to predicting the cognate epitope target of a T-cell receptor (TCR) have been proposed. However several questions on the advantages and disadvantages of these different approaches remain unresolved, as most methods have only been evaluated within the context of their initial publications and data sets. Here, we report the findings of the first public TCR-epitope prediction benchmark performed on 23 prediction models in the context of the ImmRep 2022 TCR-epitope specificity workshop. This benchmark revealed that the use of paired-chain alpha-beta, as well as CDR1/2 or V/J information, when available, improves classification obtained with CDR3 data, independent of the underlying approach. In addition, we found that straight-forward distance-based approaches can achieve a respectable performance when compared to more complex machine-learning models. Finally, we highlight the need for a truly independent follow-up benchmark and provide recommendations for the design of such a next benchmark.

## Introduction

A key challenge within immunoinformatics is the prediction of the target epitope for a T-cell receptor (TCR) sequence. Indeed, the recognition of an epitope by a T-cell receptor (TCR) is an essential step for the activation of T-cells and thus critical for a functioning adaptive immune system. Epitopes are short peptides presented by the major histocompatibility complex (MHC) on the surface of antigen-presenting cells, allowing cognate TCRs to bind them and to confer specificity to the T cell. TCRs consist of a heterodimer, most commonly of an alpha- and a beta chain. Each of these chains are the result of a V(D)J somatic recombination event during T-cell maturation. Due to the randomness of this recombination process, each T-cell clone expresses a potentially unique TCR and thus has a unique epitope specificity. This high TCR diversity allows the adaptive immune system to respond to the myriad of seen and unseen threats.

The advent of high-throughput adaptive immune receptor repertoire (AIRR) sequencing techniques allows to access the TCR sequences of large parts of T-cell repertoires. However, the sheer numbers of existing TCRs mean that most of the TCRs encountered in an experiment may not have been characterized before. Moreover, while the presence of specific TCRs in a population of T-cells can now often be established, their cognate epitope targets mostly remain unknown. Because epitope recognition is crucial for pathogen defense, vaccine response, tumor control and autoimmune diseases and since TCR specificity helps understanding the function of a T cell, it is essential to learn to decipher it.

During the past years, several solutions to unravel and predict the specificity and the cognate epitope target of a TCR have been proposed, ranging from a simple database look-up to deep learning-based prediction models. The advantages and disadvantages of these different approaches have not yet been systematically examined, most having only been evaluated within the context of their initial publications and data sets. In addition, the annotation of TCR-epitope pairs is a complex problem: the promiscuity of TCR-epitope binding and the technical but also experimental variations underlying paired TCR-epitope data make the identification of a clear signal difficult [1]. There is thus a clear need to benchmark existing TCR-epitope prediction approaches, enabling the field to progress towards an understanding of the principles underlying T-cell specificity.

Here we report the findings of the first public TCR-epitope prediction benchmark performed in the context of the ImmRep 2022 TCR-epitope specificity workshop (https://www.pks.mpg.de/immrep22). Leading scientists in the field as well as junior researchers interested in the TCR specificity problem were invited to participate and were offered datasets to train and test a collection of existing prediction models. The aim of the workshop was to evaluate and compare the obtained outputs to classify the approaches and most importantly, to help identify an ideal dataset and optimal evaluation strategies for future follow-up efforts. Describing the outcome of the workshop, we attempt to group the selected and tested methods, both - based on the TCR feature input as well as on the underlying prediction algorithm - in an attempt to identify patterns within the performance results. We conclude the report with lessons learned on the TCR-epitope problem from this first benchmarking study and make several recommendations towards future attempts at benchmarking.

## Materials and Methods

### Construction of the training- and test data

The benchmark data set was derived from the VDJdb database (downloaded on 23/06/2022). VDJdb is a curated database of TCRs with known antigen specificities [2]. Only those TCR-epitope pairs with paired alpha-beta chain data were collected. Any duplicated TCR-epitopes were removed, as defined by V/J gene usage as well as the CDR3 (complementarity determining region) sequence for both the alpha- and the beta chain. Only TCR-epitope pairs obtained by tetramer- or dextramer-sort were retained. Furthermore, all the dextramer-sort entries originating from the 10X technical report [3] were excluded because of the high reported cross-reactivity of these TCRs [1]. Lastly, only those 17 MHC-epitopes that had at least 50 unique TCR sequences were retained.

A full list of epitopes and the number of their associated TCR is provided in Table S1. To constitute the negative control data set, unpublished paired alpha-beta chain TCR sequences without peptide specificity information and obtained from 10x genomics sequencing of CD8+CD96+ T cells from 11 control individuals were provided by A. Eugster. The entire data was subsequently separated into “positive”/”negative” training- and test data sets as follows.

The “positive” set for each epitope under consideration was extracted from the VDJdb as described. The “negative” set for each epitope was constructed by randomly sampling a set of TCRs specific to any of the other 16 epitopes. The size of this negative set was three times larger than the number of the positive TCRs for the epitope in consideration. The negative set was further expanded by randomly sampling TCRs from the negative control dataset to obtain twice the number of positive TCRs. Thus, the final set for each epitope had a negative/positive ratio of 5:1. For instance, for the epitope “ATDALMTGF” with 132 positive TCRs, there are 660 negative TCRs, of which 396 TCRs originated from the swapping of the TCRs from other epitopes while 264 TCRs were sampled from the negative control data. The data was then split randomly into a training and test set in the ratio of 80:20.

### Models applied

In total, 23 TCR-epitope prediction models were trained and tested during the course of the workshop, which can be found in table 1. Most approaches have been previously published, or are novel variants of existing models. These variations were created specifically for the workshop to explore the added benefit of integrating specific information types for the TCR-epitope prediction problem, notably the integration of specific TCR chain data. For comparison purposes, a ‘Random’ model was included, which produced scores between zero and one, based on the Numpy random number generator.

**Table 1:**
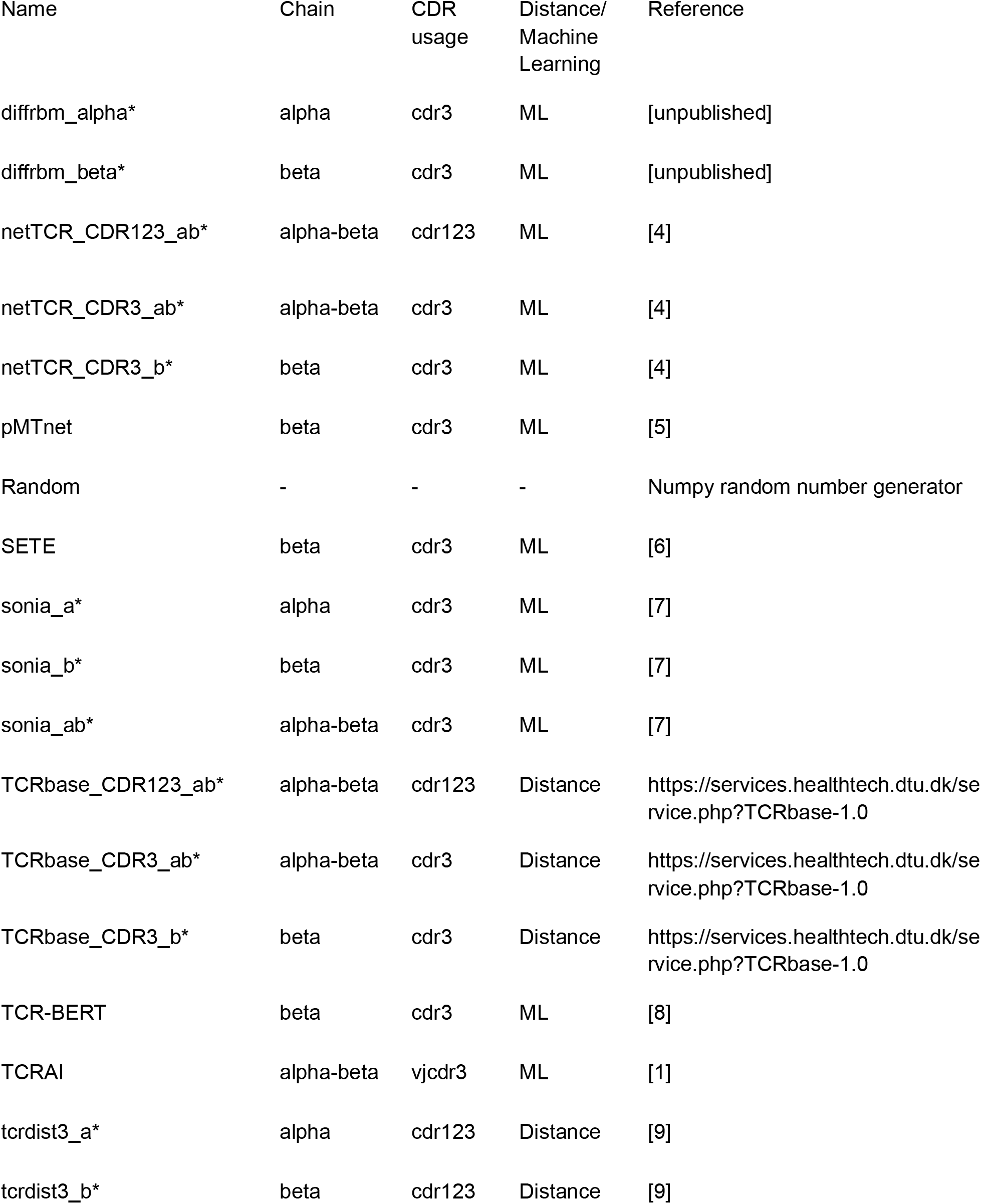
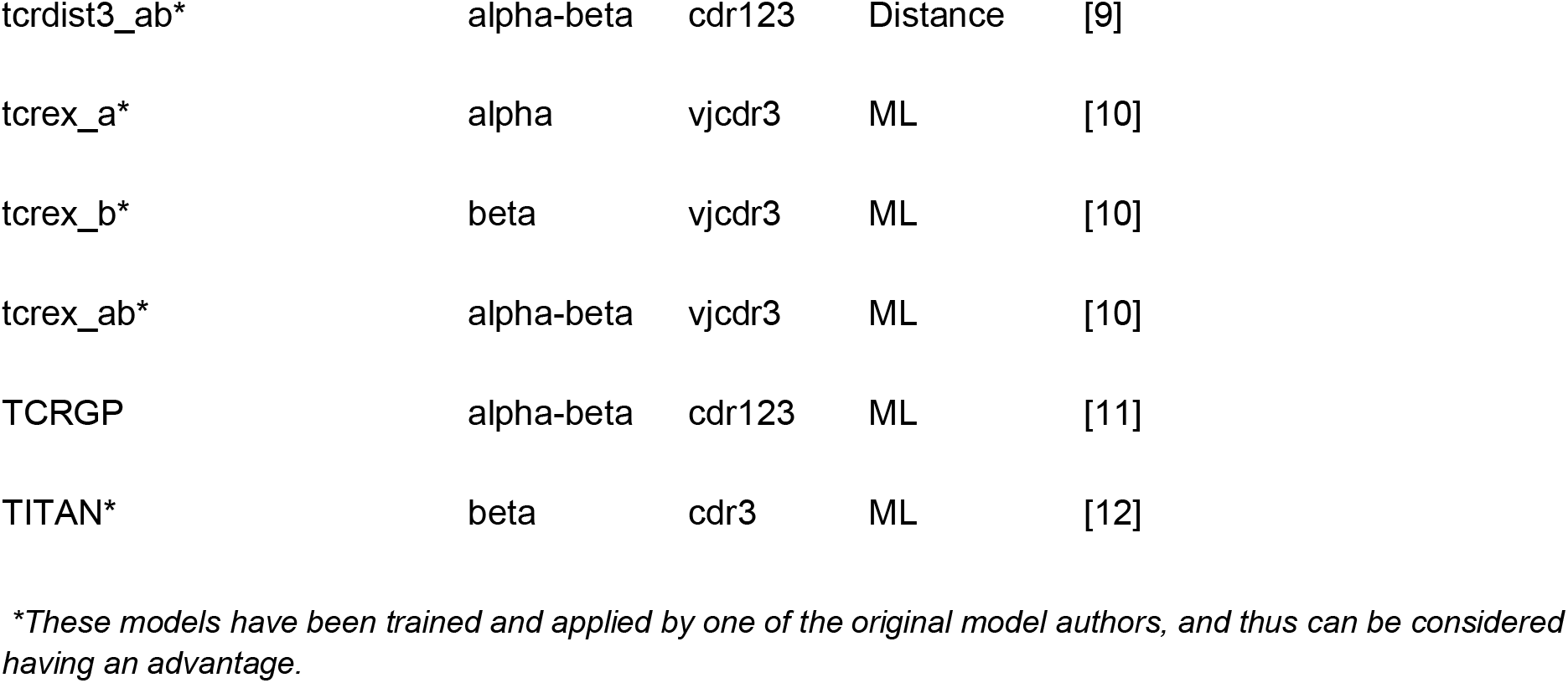
List of models tested.

### Evaluation of prediction performance

Models were trained on the available training set for each epitope, or on all available training sets where appropriate for multi-label models. To decrease information leakage, the models were only allowed to train on the available data and any pre-training steps on TCR specificity were excluded. A decision value was then assigned to each TCR in the test data set for a given epitope with a blinded ground truth with respect to negative and positive samples. The ground truth of the test set was thus not available to workshop participants during model training and application.

Two data set setups were utilized to evaluate the models. The first included a mixture of positive and negative test data for each epitope, with the target epitope being known. Thus, each model had to score the likelihood of the TCRs included in the dataset binding to the specified epitope. From these results, the area under the ROC curve (AUC) was calculated. In the second setup, all positive test data from the previous setup were merged, and each method was challenged to provide predictions for every possible epitope seen during training. The epitopes were then ranked for each TCR from the most likely to the least likely according to the predictions of the model, and the rank of the true epitope was enumerated. As an epitope has multiple true TCRs, an average rank was calculated across all TCRs for one epitope. All prediction scores were then collected and analyzed with the same evaluation script, which calculated the AUC / the average rank for each epitope, as well as the average over all epitopes.

### Github and data repository

The data sets and evaluation scripts can be found at https://github.com/viragbioinfo/IMMREP_2022_TCRSpecificity.

## Results

### No relation between size of the training data and model performance

When benchmarked on the 17 MHC-epitope test data, all methods reached a non-random performance (AUC > 0.5) for most epitopes, as can be seen in figure S1. This demonstrates that independent of the method used, it is possible to classify unseen TCRs for a given epitope within this dataset. Only the SARS-CoV-2-derived epitope NQKLIANQF consistently scored poorly or even randomly across all methods (AUC 0.539 mean ± 0.065 s.d.). In contrast, the easiest to predict epitope, NYNYLYRLF, also a known SARS-CoV-2 epitope, featured a near-perfect classification for most methods (AUC 0.956 mean ± 0.042 s.d.). TCRs for both the NQK and NYN epitopes were derived from the same MHC-dextramer study [13], thus there is no experimental difference in how the TCRs were collected or from which individuals. Furthermore, both epitopes had very similar training data sizes, namely 112 and 88 respectively. An overall analysis did show a strong relationship the performance and the similarity between TCR sequences in the training set (quantified as the average Levenshtein distance between the CDR3 sequences), as can be seen in Figure 1. The strongly performing NYN epitope had an average Levenshtein distance of 12.6 within its training CDR3 sequences, while the NQK epitope featured an average distance of 17.7. This relationship was found to be consistent across most tested methods, as can be seen in figure S4. Therefore, those epitopes that had many highly similar TCR sequences seemed to be easier to classify within the presented held-out data set-up.

**Figure 1:**
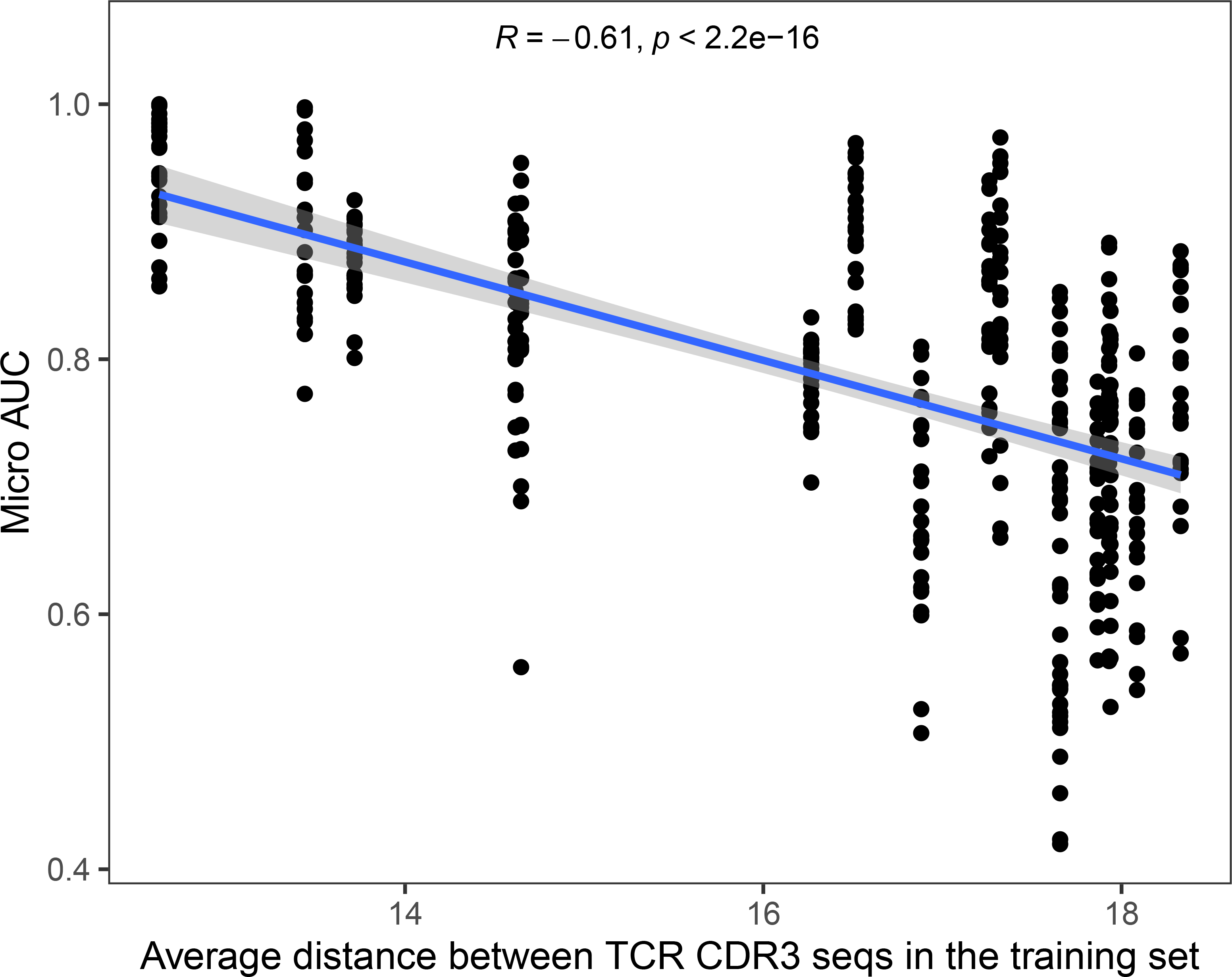
Relationship between the prediction performance and similarity of TCR sequences in the training set. The x-axis shows the average Levenshtein distance between the CDR3 sequences in the training data set. The y-axis shows the micro AUC for each epitope and each method.

### Distance-based methods provide a good baseline prediction

Methods that annotate unseen TCRs for binding a specific seen epitope can be broadly divided into two categories, namely distance-based methods and feature-based classification methods. Distance-based methods, such as TCRbase, mainly use a single distance metric to calculate the similarity between unseen TCRs and seen TCRs in the training data, independent of the epitope. In its simplest iteration, this can be the amount of amino acid mismatches between the two CDR3 sequences (i.e. the Hamming distance). If the distance is below a given threshold, an unseen TCR can be judged sufficiently similar to the training set TCR to be annotated with the same epitope. The distance metric itself can then be considered a confidence estimate of the method, where larger distances are considered less reliable annotations. More commonly, these distance-based methods rely on a k-nearest neighbor approach (k-NN), as is the case for all distance-based methods used in this benchmark. Within these k-NNs, it is the label of the k most similar TCRs that is used to predict the target epitope.

Feature-based methods are defined here as those that try to identify common patterns underlying the training set TCRs that bind a given epitope. These patterns then form the basis of predicting the binding preference of unseen TCRs. The underlying model is always a supervised machine learning method, where the known TCRs binding each epitope are provided as training data. The result is therefore a fitted model, which can then be applied to any unseen TCR. These are distinct from the distance-based methods as they try to learn which features are important for each epitope. However, as can be seen in figure 2, some distance-based approaches have a performance that is very close to those of the best performing feature-based methods. This supports the use of these methods as a comparative baseline, as any new, more complex methodologies claiming to learn TCR-epitope patterns should be required to outperform basic matching algorithms.

**Figure 2:**
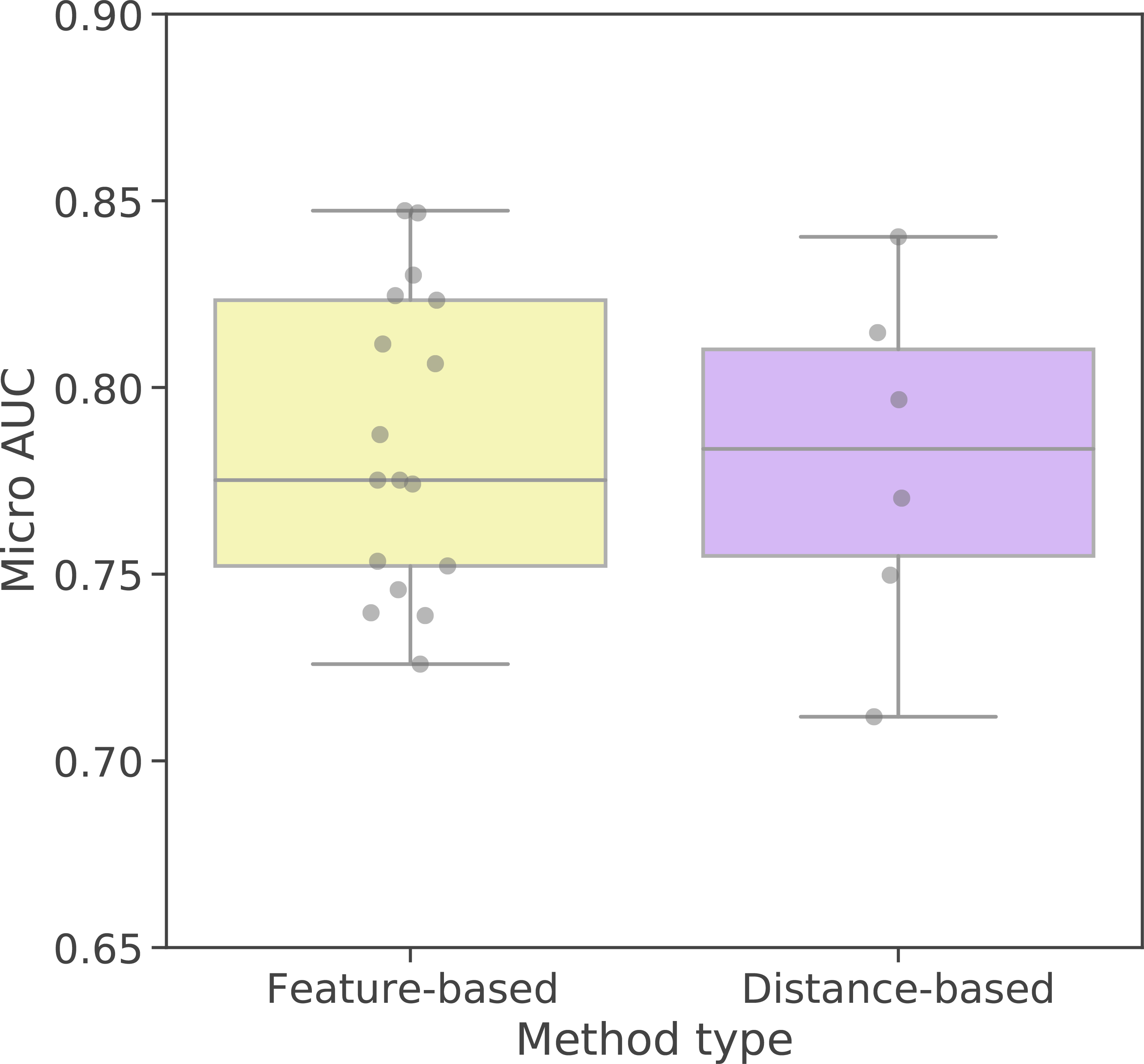
Average microAUC by approach. Distance-based methods to the left annotate TCRs based on the similarity to known epitope-specific TCRs. Feature-based approaches use machine learning to learn associated features specific to an epitope, which is then used for classification.

An additional distinction can be made between those machine-learning approaches that train one model for each epitope separately (peptide-specific) or one model for all epitopes simultaneously (pan-specific). However, research has shown that even in the latter case, most pan-specific methods internally act as having a model per epitope [12]. This is because the generalisation of epitope-TCR pairing across different epitopes is still currently challenging due to the so far rather limited number of peptides characterized by TCR data [14]. This also matches with the found results in this study, as there seemed to be no consistent difference between these two approaches, as can be seen in Table S2.

### Combining alpha and beta chain improves epitope prediction

Historically, the majority of methods to address the TCR-epitope specificity problem focus on the CDR3 sequence of the beta chain only for predictions, as historical TCR sequencing efforts have mainly only characterized the TCR beta chain. However, the TCR heterodimer complex exists of both alpha and beta chains, and both are known to make contacts with the epitope [15]. Paired alpha-beta chain information is still rare as it requires TCR sequencing at the single cell level, but prior studies have suggested that prediction performance can be increased by using information from both chains, alpha and beta [16]. The current benchmark data set only included the TCR-epitope entries with paired alpha- and beta chain information. This allowed a direct comparison between methods only using one of the two chains, by only using the relevant part of the input data. In this manner, the difference in prediction performance using either alpha, beta, or both chains could be assessed on the same TCR-epitope pair training- and test data. As can be seen in figure 3, methods that use both chains (alpha and beta) consistently outperform methods that only use a single chain (alpha or beta). The same trend can be seen at the level of individual AUCs, as seen in table S2 and figure S2, where the most performant method per epitope is usually one that uses both chains. Furthermore, when clustering the individual performances, the use of single or both chains can be seen as the most prominent grouping factor, as can be seen in figure S3.

**Figure 3:**
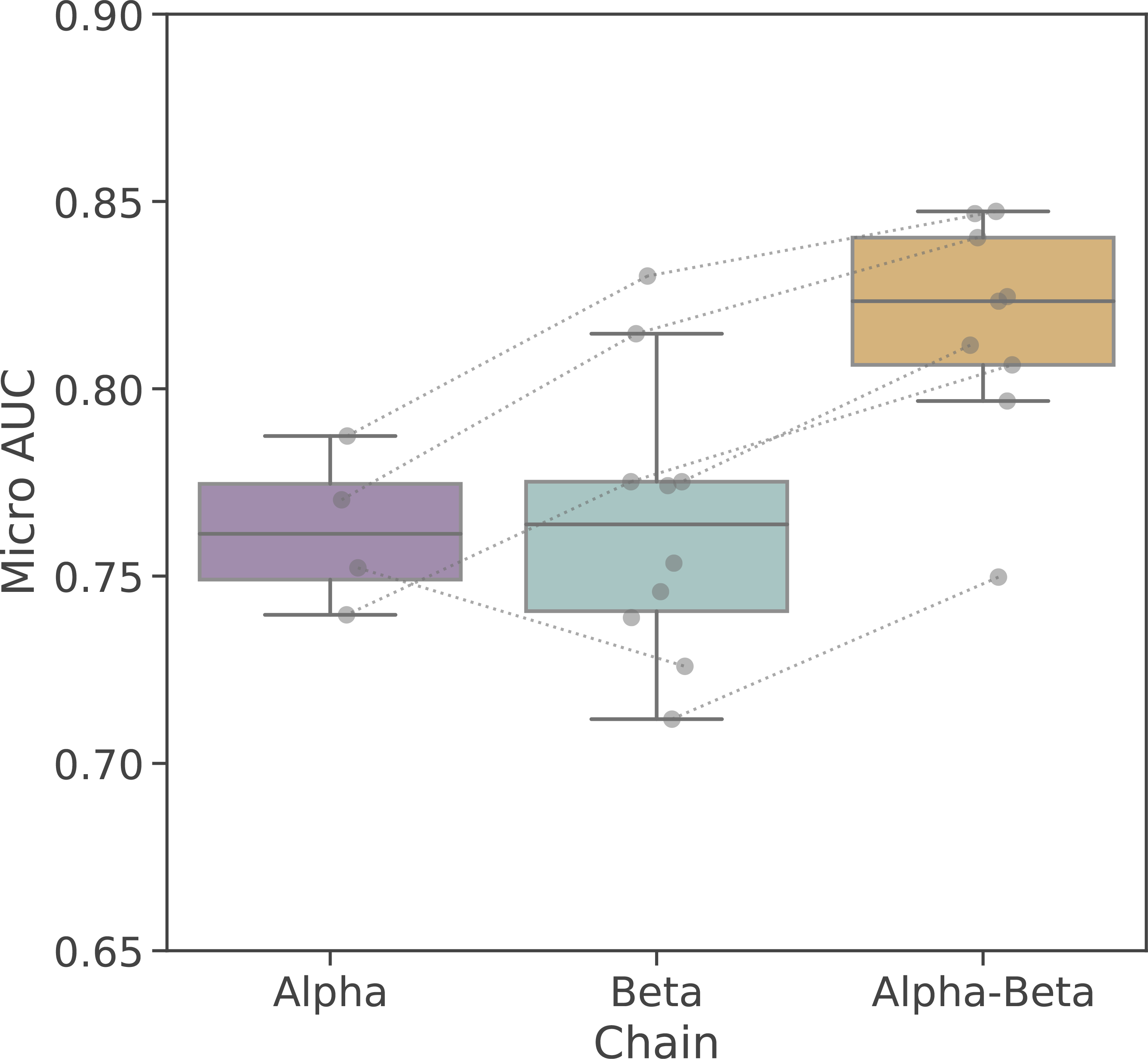
Comparison of average microAUC of methods considering both TCR chains (Alpha-Beta) and methods that only consider the beta chain (Beta) or the alpha chain (Alpha). The lines denote those methods that use the same architecture but with different input.

### Integrating V/J gene usage or CDR1/CDR2 improves epitope prediction

Most methods focus only on the CDR3 region of the beta (or alpha chain) of the TCR, as this region is the most variable and responsible for the majority of contacts with the epitope residues. Even though the CDR1 and CDR2 of a TCR are wholly determined by the V gene usage, they add a degree of variability to the chains and facilitate the crucial contacts between the TCR and the epitope-MHC complex. To investigate the complementary information contained in V/J genes and CDR1/CDR2 segments, we can divide methods into three categories based on their inputs: i) Only requiring CDR3 amino acid sequence (cdr3) as the input, ii) Requiring the CDR1/CDR2/CDR3 amino acid sequence (cdr123) as the input, and iii) Requiring the CDR3 amino acid sequence as well as the V/J gene usage (vjcdr3). From the average performance results, as seen in figure 4, we observed that methods that consider the CDR1/CDR2 regions, either directly (cdr123) or indirectly (vjcdr3), outperform those that only consider the CDR3 amino acid sequence. However, considering the amino acid sequence of the CDR1 and CDR2 regions does not, in this setting, seem to have any improvement over simply considering the V gene annotation itself. Some methods, such as tcrdist3, do utilize a prior alignment of these regions to calculate their difference (as well as an additional variable loop between CDR2 and CDR3 which is given the name “CDR2.5”), and thus already formulate the concept that some CDR1/CDR2 are more similar than others.

**Figure 4:**
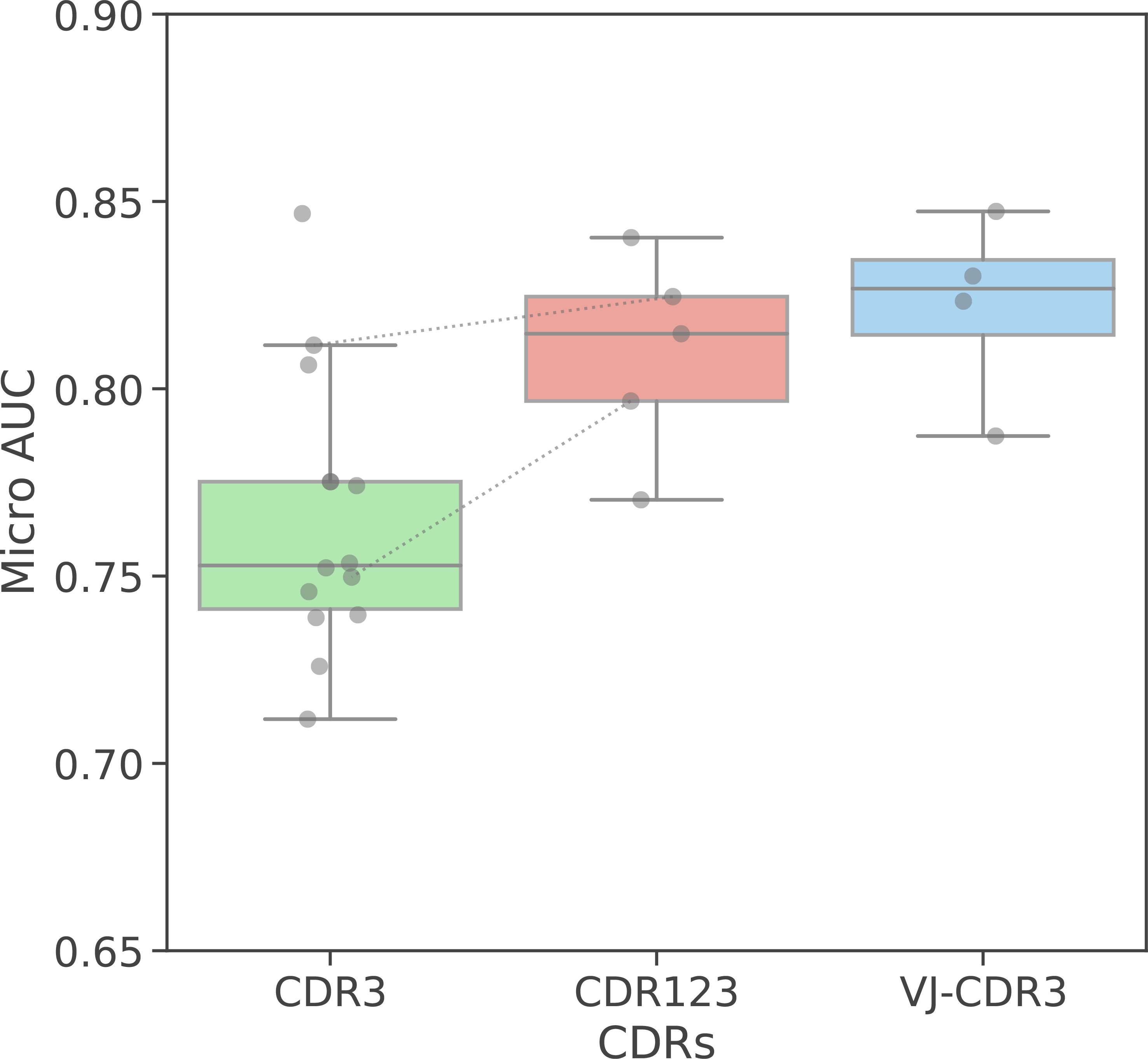
Average performance by CDR region usage. From left to right, methods using only CDR3 amino acid sequence, methods that use CDR3 and V/J genes, and methods that use CDR1,CDR2,CDR3 amino acid sequences as input. The lines denote those methods that use the same architecture but with different input.

### Epitope ranking mostly, but not always follows binary classification performance

Prior sections have focused on method performance as calculated by the AUC of the ROC curve, thus the trade-off between false negatives and false positives for the binary classification problem. However, epitope-TCR pairing is not a true binary classification problem, as each TCR needs to be theoretically matched with any epitope. For this reason, we considered the epitope rank as an alternative metric, where the score for the correct epitope of a TCR is judged against all other trained epitopes on the same TCR. In this instance, a lower score is considered better and a score of 1 signifies a perfect prediction for an epitope. As this required applying the model for each epitope on the same data set, this could not be accomplished for every method previously tested. Some methods were not able to create models for each epitope, others did not provide the option to predict per epitope. As can be seen in Figure 5, the relative performance across models between AUC and ranked epitope has remained relatively stable. The most performant method with regards to AUC remains the most performant method with regards to the epitope rank. This is not unsurprising as the AUC already captures the difference in prediction scores between a positive pair and a negative pair. The key difference is that the epitope rank focuses solely on the positive pairs. Indeed, the overall ranking between some methods does change, as can be seen in supplemental table S3. Most notably, basic clustering approaches, such as TCRbase, score relatively better on average epitope rank than AUC. This can be attributed to the fact that these methods do not hold the concept of a negative background, as they are searching for TCRs similar to those TCRs known to bind an epitope. Thus, it can be expected that these methods are less able to distinguish negative samples, despite being proficient in the annotation of positive samples with the right epitope.

**Figure 5:**
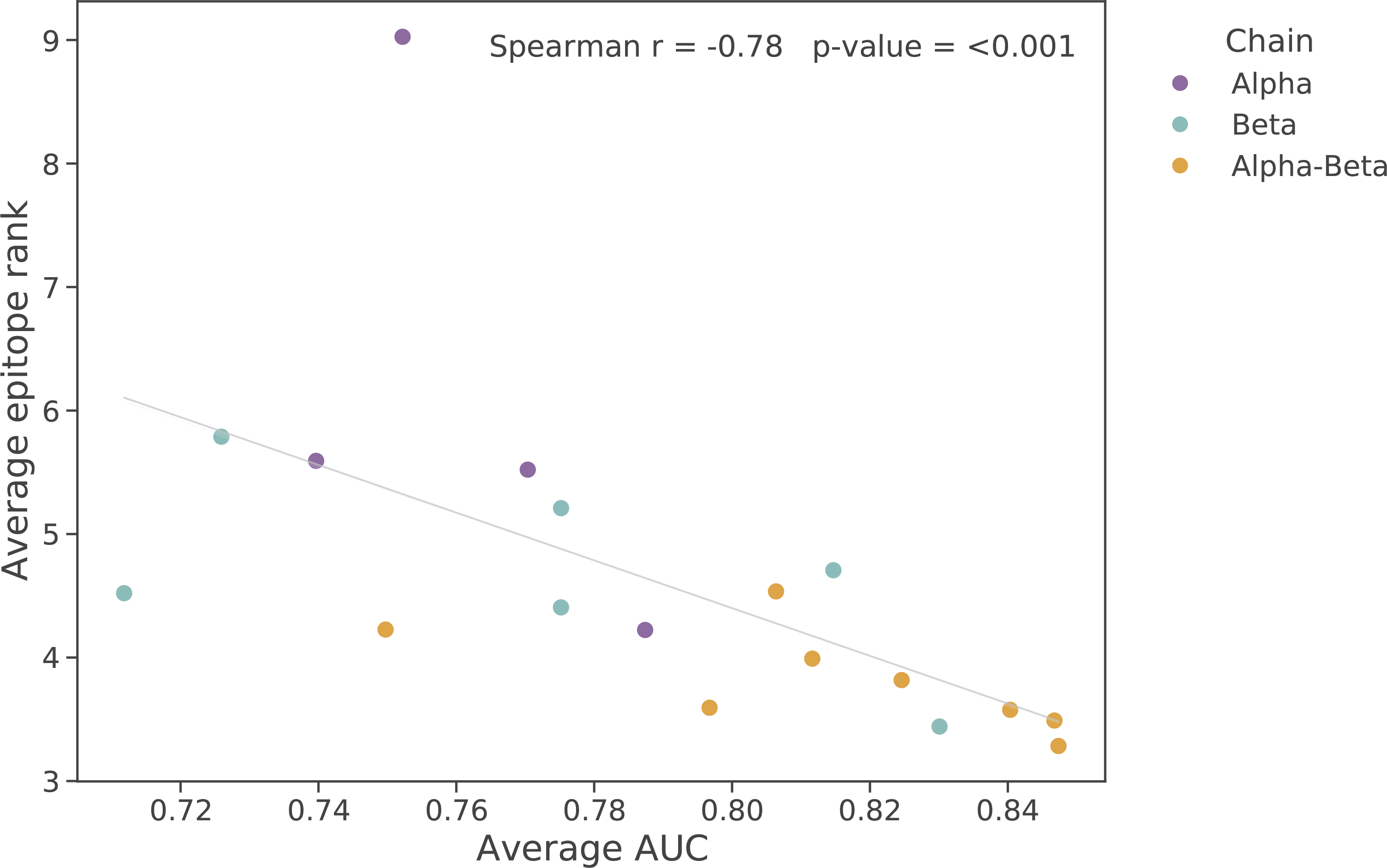
Average performance of methods based on microAUC in the binary classification problem (x-axis) plotted against the epitope rank performance for multi-label prediction (y-axis). Each dot in the plot represents a single method.

## Discussion

### Limitations of the benchmark

While many methods were included into this benchmark, our effort was not exhaustive. The wealth of currently existing methods for TCR-epitope prediction makes it impossible to compare them all in a single effort. In addition, many methods that have been described in the literature unfortunately lack a publicly accessible code/interface.

In addition, ground-truth true negative data does not exist within the TCR-epitope context. Two strategies to circumvent this problem were combined for this benchmark: swapped TCR-epitope pairs and unrelated repertoires from healthy individuals without epitope knowledge as background sequences. Both have their advantages and disadvantages. Swapping TCR-epitope pairs by considering the TCRs positive for one epitope as being negative for another, means that methods can in theory learn the TCR patterns associated with the negative epitopes in training and test data. Utilizing unrelated, healthy repertoires has the disadvantage that there may be an experimental and/or biological bias in the data. This lends itself to the possibility that any model may learn the distinction between positive and negative based on such biases alone. In addition, healthy repertoire data is not free of potential misannotations, due to the presence of T cells specific for common dominant epitopes also present in the positive data. As highlighted by the difference in performance ranking based on AUC and epitope ranking, classifiers potentially do have a tendency to learn patterns within the negative data that changes how they will consider the problem. This benchmark is far too limited to provide a solid answer to what negative strategy is most appropriate, and this remains an important topic for future research.

Another key choice involved the lack of removal of similar TCRs from the test set compared to the training data set. This would have required a strict definition of what constitutes a sufficiently similar TCR. There does not currently exist a common consensus of how different a TCR can be before it can be expected to no longer bind the same epitope. Furthermore, whether to include or exclude similar TCRs from the datasets is a decision that depends fully on the downstream application. Mostly, the end goal of these methods is to identify epitope-specific TCR pairs among a large TCR sequence repertoire extracted independently. It is already known that public clonotypes do occur across different individuals, and that small subsets of TCRs can be successfully annotated with their epitopes just by matching their TCR sequences to annotated TCR sequences of a database [17–19]. However, public clonotypes are the exception rather than the rule. Removal of the highly similar TCRs from the test set does allow evaluation to what degree these methods are capable of linking distant TCRs to the correct epitope. This results in a far more limited positive test set, with more focus on the classification of the negative samples.

Finally, many methods are hampered in their success by the current limited training data set size. Thus their measured performance is not representative of their theoretical potential with an unrestricted training data set. For example, several methods have shown improvements by utilizing transfer learning or pre-training steps, which are not possible with this limited benchmark [5]. In addition, methods using only beta-chain data can typically rely on a larger dataset than methods with both alpha and beta chains, which may explain their poorer performance in this instance. The performing data set curation from paired chain data also had an impact on the size of the training and the test set of a few epitopes. For instance, two TCRs would be considered for evaluation if their alpha chains are different, implying that they are overall different but they may collapse to a single TCR if their beta chain is identical. Furthermore, the dataset used here and split into test- and training data sets was derived from the same experiments. This was necessary given the current scarcity of available TCR-epitope specific data. Ideally, the train-test split would be derived from independent experiments to avoid any experimental information bleed, however this can only be done for a very limited number of epitopes.

### The shape and scope of future benchmarks

The goal of this study was to evaluate the possibility of a benchmark between methods and highlight important lessons learned from this pilot study. Due to the aforementioned limitations, no strong conclusions should be made about the superiority of one approach to another at this point. Regardless of the limitations, an independent benchmark has become necessary due to a rapid surge in the number of the TCR-epitope prediction tools. It is no longer feasible or reasonable to expect a single study to compare to all other existing methods. Furthermore, as highlighted in this study, the choice of data set on which to evaluate can have a large impact on the performance. Related is the commonly accepted phenomenon that a method will always score best when applied by its own authors and on the data set in the paper where it is introduced.

Other prediction-focused fields have embraced the idea of an open competition where methods are benchmarked on the same never-before-seen data set. This is a way to independently evaluate the wide variety of methods that are available, but also to drive the field forward and enable it to reach new heights. As a conclusion of the ImmRep TCR-epitope specificity workshop, we strongly encourage the organization of similar competitions focused on the TCR-epitope prediction problem. The most essential choice will be the origin of the test data as many options are available, from a stratified approach as highlighted here, to the integration of simulated datasets with a known ground truth [20]. The ideal data set for such a challenge would unequivocally be an unpublished independent data set with both TCRs with known epitope-specificity, as well as TCRs without this information.. As highlighted in this study, it would ideally involve paired alpha-beta TCR sequence data, and would therefore likely be derived from a single cell sequencing experiment. In addition, the use of oligo-tagged multimers would enable both identification of those TCRs that are specific for an epitope, along with those that are likely not. Furthermore, this data set should contain multiple previously studied epitopes, so as to compare the epitope rank beyond the straight-forward classification problem. The technology to create such a dataset is currently available, and therefore only requires the willingness of either funders, institutes or companies to provide it.

## Conclusions

This study contains an initial large-scale benchmark of epitope-TCR prediction methods. It was not meant to be exhaustive, nor should the results be overly interpreted as the data set was limited and the evaluation superficial. Several important observations could nevertheless be established. The use of paired-chain alpha-beta data, as well as CDR1/2 or V/J information, improves classification when this data is available, independent of the underlying approach. Straight-forward clustering approaches can achieve a respectable performance and should be used as a valid benchmark for future studies. Finally, there is a large need for a true independent benchmark on the myriad of methods within the field.

## Acknowledgements

We wish to thank the Max Planck Institute for the Physics of Complex Systems in Dresden and the Max Planck Society for their generous financial and physical support enabling the ImmRep 2022 TCR-epitope specificity workshop. We wish to thank the VDJdb resource for providing the dataset that formed the basis of this paper. In addition, we wish to thank the valuable input from Prof. Victor Greiff on the manuscript.

## Author contributions

Conceived the study: AE, VS; Analysis and data collection: PM, JB, BB, LCL, VK, EL, AM, MM, TM, PP, AP, MRM, JF, AV, AMW, AW, RY, AE, VS; Evaluation and visualisation: PM, LCL, VK; Wrote and edited the manuscript: PM, JB, BB, LCL, VK, EL, AM, MM, TM, PP, AP, MRM, JF, AV, AMW, AW, RY, AE, VS.

