## Supplemental materials for "Benchmarking solutions to the T-cell receptor epitope prediction problem: IMMREP22 workshop report"

Table S1: Epitope list

| Epitope | # Training samples |
|---|---|
| LTDEMIAQY | 200 |
| GILGFVFTL | 1088 |
| TTDPSFLGRY | 386 |
| NQKLIANQF | 112 |
| HPVTKYIM | 96 |
| GPRLGVRAT | 80 |
| KSKRTPMGF | 170 |
| CINGVCWTV | 366 |
| TPRVTGGGAM | 90 |
| SPRWYFYYL | 184 |
| LLWNGPMAV | 376 |
| GLCTLVAML | 292 |
| YLQPRTFLL | 534 |
| ATDALMTGF | 208 |
| NLVPMVATV | 548 |
| RAQAPPPSW | 72 |
| NYNYLYRLF | 88 |

Table S2 : Full list of micro AUCs for each epitope and model

| | LTDEMIAQY | GILGFVFTL | TTDPSFLGRY | NQKLIANQF | HPVTKYIM | GPRLGVRAT | KSKRTPMGF | CINGVCWTV | TPRVTGGGAM | SPRWYFYYL | LLWNGPMAV | GLCTLVAML | YLQPRTFLL | ATDALMTGF | NLVPMVATV | RAQAPPPSW | NYNYLYRLF | Average |
|---|---|---|---|---|---|---|---|---|---|---|---|---|---|---|---|---|---|---|
| **random** | 0.57 | 0.48 | 0.56 | 0.48 | 0.42 | 0.50 | 0.44 | 0.41 | 0.55 | 0.49 | 0.47 | 0.45 | 0.48 | 0.62 | 0.50 | 0.30 | 0.60 | 0.49 |
| **TCRbase_CDR3_b** | 0.63 | 0.87 | 0.60 | 0.42 | 0.78 | 0.70 | 0.82 | 0.57 | 0.56 | 0.53 | 0.81 | 0.83 | 0.77 | 0.81 | 0.55 | 0.87 | 0.98 | 0.71 |
| **diffrbm_b** | 0.63 | 0.81 | 0.65 | 0.54 | 0.76 | 0.56 | 0.80 | 0.68 | 0.84 | 0.65 | 0.73 | 0.83 | 0.70 | 0.76 | 0.58 | 0.83 | 0.97 | 0.73 |
| **SETE** | 0.56 | 0.86 | 0.53 | 0.54 | 0.69 | 0.73 | 0.92 | 0.71 | 0.69 | 0.61 | 0.77 | 0.83 | 0.74 | 0.77 | 0.64 | 0.98 | 0.99 | 0.74 |
| **sonia_a** | 0.71 | 0.86 | 0.60 | 0.61 | 0.70 | 0.85 | 0.66 | 0.76 | 0.81 | 0.59 | 0.80 | 0.82 | 0.77 | 0.72 | 0.69 | 0.77 | 0.86 | 0.74 |
| **TITAN** | 0.61 | 0.89 | 0.75 | 0.54 | 0.68 | 0.57 | 0.82 | 0.67 | 0.86 | 0.76 | 0.75 | 0.86 | 0.81 | 0.76 | 0.66 | 0.82 | 0.86 | 0.75 |
| **TCRbase_CDR3_ab** | 0.64 | 0.87 | 0.62 | 0.53 | 0.72 | 0.76 | 0.83 | 0.58 | 0.75 | 0.63 | 0.90 | 0.91 | 0.79 | 0.82 | 0.59 | 0.82 | 0.99 | 0.75 |
| **diffrbm_a** | 0.67 | 0.87 | 0.62 | 0.62 | 0.71 | 0.72 | 0.70 | 0.80 | 0.70 | 0.69 | 0.81 | 0.87 | 0.75 | 0.81 | 0.65 | 0.94 | 0.87 | 0.75 |
| **TCR-BERT** | 0.63 | 0.80 | 0.51 | 0.52 | 0.82 | 0.77 | 0.81 | 0.75 | 0.81 | 0.57 | 0.82 | 0.89 | 0.80 | 0.87 | 0.54 | 0.96 | 0.94 | 0.75 |
| **tcrdist3_a** | 0.61 | 0.86 | 0.70 | 0.56 | 0.81 | 0.75 | 0.73 | 0.84 | 0.75 | 0.67 | 0.80 | 0.89 | 0.80 | 0.82 | 0.68 | 0.92 | 0.89 | 0.77 |
| **pMTnet** | 0.67 | 0.89 | 0.66 | 0.46 | | 0.77 | 0.91 | 0.72 | 0.82 | 0.73 | 0.78 | 0.89 | 0.81 | 0.75 | 0.74 | 0.85 | 0.95 | 0.77 |
| **netTCR_CDR3_b** | 0.59 | 0.86 | 0.67 | 0.42 | 0.79 | 0.80 | 0.83 | 0.75 | 0.73 | 0.73 | 0.84 | 0.90 | 0.78 | 0.82 | 0.67 | 1.00 | 1.00 | 0.78 |
| **sonia_b** | 0.63 | 0.85 | 0.66 | 0.55 | 0.69 | 0.66 | 0.85 | 0.71 | 0.86 | 0.77 | 0.83 | 0.84 | 0.76 | 0.86 | 0.75 | 0.94 | 0.98 | 0.78 |
| **tcrex_a** | 0.69 | 0.88 | 0.63 | 0.65 | 0.70 | 0.81 | 0.67 | 0.84 | 0.84 | 0.67 | 0.81 | 0.90 | 0.80 | 0.86 | 0.74 | 0.97 | 0.91 | 0.79 |
| **TCRbase_CDR123_ab** | 0.71 | 0.89 | 0.71 | 0.51 | 0.78 | 0.80 | 0.90 | 0.72 | 0.84 | 0.76 | 0.91 | 0.96 | 0.75 | 0.89 | 0.62 | 0.83 | 0.97 | 0.80 |
| **sonia_ab** | 0.71 | 0.88 | 0.68 | 0.62 | 0.75 | 0.78 | 0.88 | 0.82 | 0.85 | 0.75 | 0.89 | | 0.78 | 0.90 | 0.77 | 0.84 | 0.99 | 0.81 |
| **netTCR_CDR3_ab** | 0.75 | 0.88 | 0.66 | 0.49 | 0.80 | 0.82 | 0.87 | 0.87 | 0.81 | 0.65 | 0.91 | 0.92 | 0.82 | 0.86 | 0.69 | 1.00 | 1.00 | 0.81 |
| **tcrdist3_b** | 0.71 | 0.91 | 0.77 | 0.52 | 0.75 | 0.85 | 0.95 | 0.77 | 0.94 | 0.78 | 0.85 | 0.92 | 0.80 | 0.86 | 0.70 | 0.87 | 0.91 | 0.81 |
| **TCRAI** | 0.74 | 0.91 | 0.75 | 0.58 | 0.80 | 0.89 | 0.85 | | 0.90 | 0.71 | 0.88 | 0.93 | 0.81 | | 0.77 | 0.84 | 0.98 | 0.82 |
| **netTCR_CDR123_ab** | 0.78 | 0.91 | 0.81 | 0.52 | 0.75 | 0.73 | 0.88 | 0.88 | 0.92 | 0.75 | 0.92 | 0.95 | 0.79 | 0.87 | 0.73 | 0.91 | 0.92 | 0.82 |
| **tcrex_b** | 0.67 | 0.90 | 0.77 | 0.54 | 0.85 | 0.66 | 0.97 | 0.80 | 0.95 | 0.84 | 0.86 | 0.95 | 0.81 | 0.91 | 0.77 | 0.88 | 0.97 | 0.83 |
| **tcrdist3_ab** | 0.77 | 0.90 | 0.74 | 0.54 | 0.84 | 0.89 | 0.95 | 0.86 | 0.84 | 0.82 | 0.89 | 0.97 | 0.80 | 0.94 | 0.74 | 0.87 | 0.93 | 0.84 |
| **TCRGP** | 0.76 | 0.92 | 0.80 | 0.62 | 0.76 | 0.86 | 0.90 | 0.87 | 0.89 | 0.82 | 0.90 | 0.94 | 0.81 | 0.89 | 0.80 | 0.90 | 0.94 | 0.85 |
| **tcrex_ab** | 0.72 | 0.90 | 0.79 | 0.65 | 0.85 | 0.72 | 0.96 | 0.87 | 0.86 | 0.81 | 0.86 | 0.96 | 0.83 | 0.93 | 0.77 | 0.91 | 0.99 | 0.85 |

Table S3: Full list of average ranks for each epitope and model

| | LTDEMIAQY | GILGFVFTL | TTDPSFLGRY | NQKLIANQF | HPVTKYIM | GPRLGVRAT | KSKRTPMGF | CINGVCWTV | TPRVTGGGAM | SPRWYFYYL | LLWNGPMAV | GLCTLVAML | YLQPRTFLL | ATDALMTGF | NLVPMVATV | RAQAPPPSW | NYNYLYRLF | Average |
|---|---|---|---|---|---|---|---|---|---|---|---|---|---|---|---|---|---|---|
| **tcrex_ab** | 3.74 | 3.30 | 3.06 | 5.07 | 3.21 | 3.86 | 1.79 | 2.85 | 6.13 | 2.73 | 3.82 | 1.73 | 4.31 | 2.44 | 3.12 | 3.61 | 1.08 | 3.28 |
| **tcrex_b** | 4.72 | 3.34 | 3.63 | 5.87 | 3.29 | 5.23 | 1.50 | 3.91 | 2.79 | 2.63 | 2.82 | 2.07 | 4.66 | 2.87 | 3.51 | 4.00 | 1.67 | 3.44 |
| **TCRGP** | 4.04 | 3.10 | 3.43 | 5.33 | 4.92 | 2.36 | 2.88 | 3.43 | 4.25 | 3.75 | 2.79 | 2.49 | 4.04 | 3.31 | 3.43 | 3.44 | 2.33 | 3.49 |
| **tcrdist3_ab** | 5.22 | 2.63 | 4.46 | 6.57 | 4.04 | 3.23 | 2.02 | 3.47 | 4.42 | 4.06 | 2.51 | 1.74 | 4.32 | 2.40 | 4.41 | 3.22 | 2.08 | 3.58 |
| **TCRbase_CDR123_ab** | 5.44 | 1.68 | 3.39 | 8.80 | 5.67 | 5.73 | 2.17 | 2.46 | 4.08 | 4.00 | 2.21 | 1.73 | 2.96 | 2.46 | 2.94 | 3.56 | 1.83 | 3.59 |
| **netTCR_CDR123_ab** | 3.40 | 3.97 | 3.43 | 5.00 | 4.25 | 4.18 | 3.55 | 3.39 | 3.42 | 4.92 | 2.98 | 3.16 | 5.69 | 4.19 | 2.65 | 3.89 | 2.83 | 3.82 |
| **netTCR_CDR3_ab** | 4.00 | 4.17 | 4.94 | 4.00 | 3.92 | 3.09 | 3.79 | 4.17 | 4.42 | 5.79 | 3.19 | 3.89 | 5.42 | 5.46 | 3.49 | 3.11 | 1.00 | 3.99 |
| **tcrex_a** | 4.16 | 3.60 | 4.94 | 4.73 | 5.71 | 4.18 | 5.38 | 3.54 | 6.25 | 4.02 | 4.68 | 2.89 | 4.60 | 3.87 | 4.45 | 2.00 | 2.79 | 4.22 |
| **TCRbase_CDR3_ab** | 6.28 | 1.68 | 3.92 | 8.33 | 6.50 | 6.55 | 3.17 | 4.02 | 5.83 | 5.63 | 2.32 | 2.11 | 2.93 | 3.38 | 3.65 | 3.89 | 1.67 | 4.23 |
| **netTCR_CDR3_b** | 6.20 | 4.46 | 5.35 | 6.60 | 3.75 | 2.91 | 3.50 | 6.04 | 3.25 | 5.25 | 3.51 | 4.49 | 6.24 | 5.35 | 3.57 | 3.44 | 1.00 | 4.41 |
| **TCRbase_CDR3_b** | 6.00 | 1.74 | 4.16 | 10.60 | 6.08 | 8.45 | 2.74 | 4.05 | 6.92 | 4.63 | 2.51 | 2.59 | 3.51 | 3.54 | 3.72 | 4.11 | 1.50 | 4.52 |
| **sonia_ab** | 5.12 | 2.03 | 3.73 | 8.53 | 8.75 | 9.09 | 3.79 | 3.02 | 6.00 | 4.75 | 2.55 | 2.89 | 3.07 | 3.96 | 2.09 | 4.89 | 2.83 | 4.54 |
| **tcrdist3_b** | 7.32 | 2.17 | 5.13 | 8.43 | 6.50 | 6.14 | 2.40 | 5.18 | 4.21 | 5.60 | 3.87 | 2.74 | 4.43 | 4.42 | 5.21 | 3.00 | 3.25 | 4.71 |
| **TITAN** | 5.88 | 3.88 | 4.20 | 7.33 | 8.33 | 5.73 | 5.12 | 5.61 | 3.92 | 2.46 | 5.32 | 3.32 | 4.82 | 4.73 | 5.38 | 3.67 | 2.00 | 4.81 |
| **sonia_b** | 5.84 | 2.35 | 3.67 | 11.00 | 7.58 | 10.82 | 4.55 | 4.00 | 9.25 | 4.88 | 2.53 | 3.78 | 3.52 | 4.38 | 2.29 | 5.78 | 2.33 | 5.21 |
| **tcrdist3_a** | 7.42 | 2.71 | 5.72 | 8.07 | 7.17 | 7.73 | 7.07 | 4.46 | 5.75 | 7.92 | 4.47 | 3.03 | 4.28 | 5.17 | 5.40 | 3.06 | 4.46 | 5.52 |
| **sonia_a** | 5.20 | 1.96 | 4.22 | 8.53 | 9.67 | 8.82 | 8.02 | 3.89 | 7.75 | 6.79 | 4.21 | 3.57 | 3.03 | 6.23 | 2.93 | 5.67 | 4.58 | 5.59 |
| **random** | 8.60 | 8.76 | 8.63 | 8.67 | 8.67 | 8.55 | 11.07 | 8.83 | 9.50 | 8.13 | 8.94 | 8.24 | 9.03 | 9.31 | 9.64 | 10.22 | 10.33 | 9.12 |

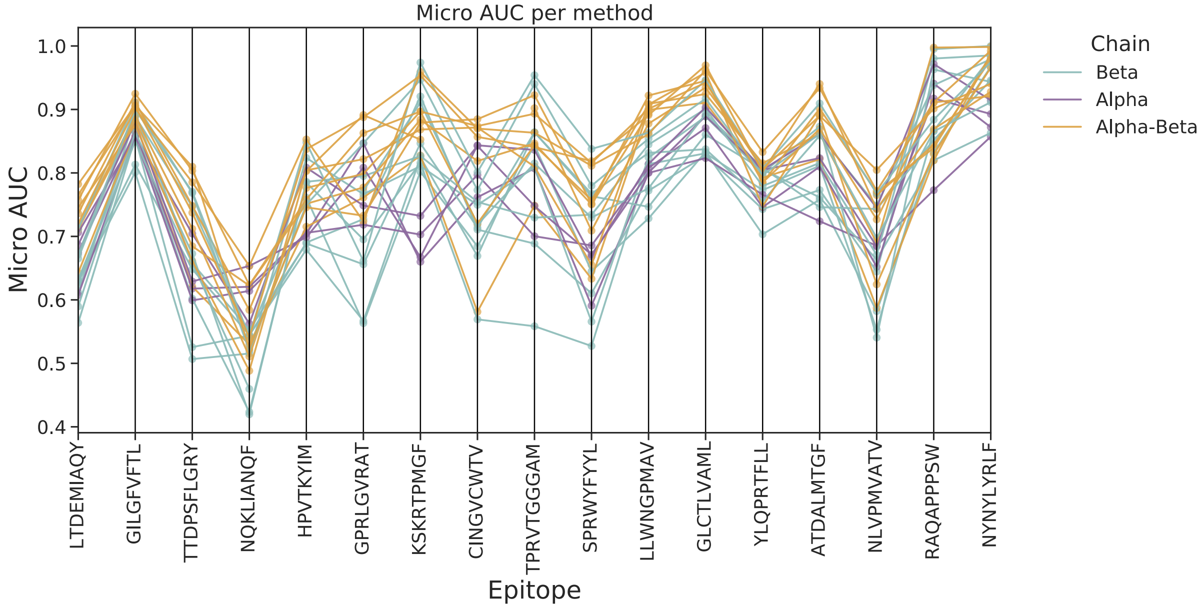


**Figure S1: Detailed MicroAUCs per epitope.** On the x-axis, all tested epitopes are listed. On the y-axis is the micro AUC as measured for each method. The line colors denote if the method used alpha-chain, beta-chain, or both as input data. Notably for most epitopes, the best performing methods were those that used both chains.

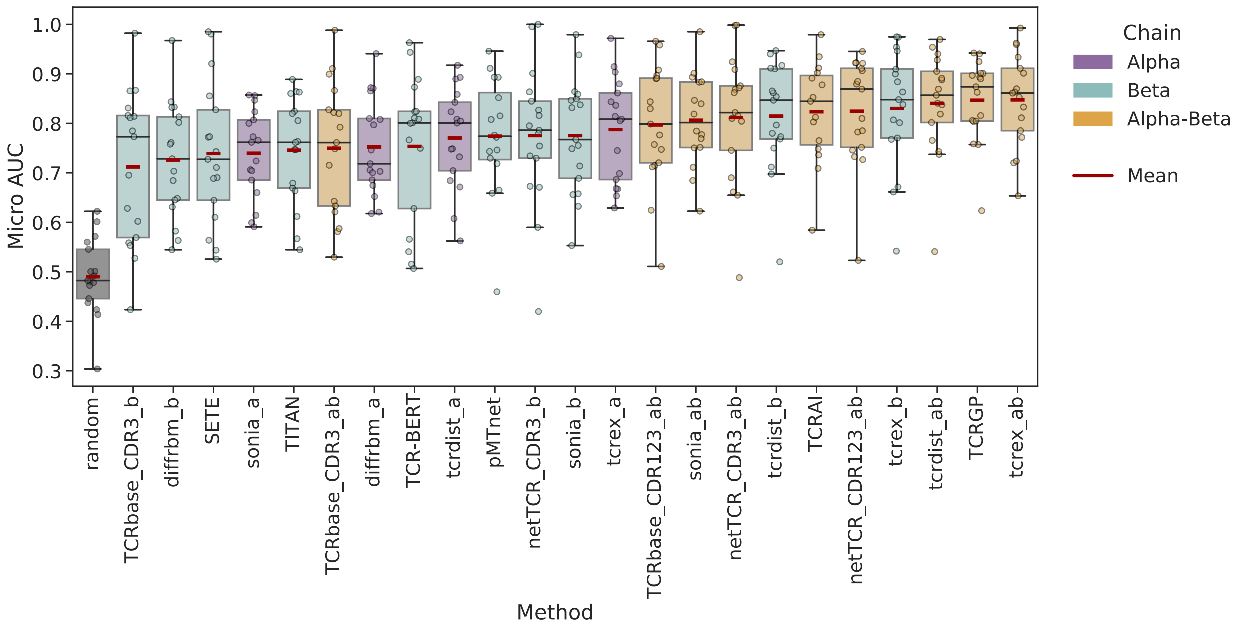


**Figure S2: Detailed MicroAUCs per method.** On the x-axis, all tested models are listed. On the y-axis is the micro AUC as measured for each epitope. The box colors denote if the method used alpha-chain, beta-chain, or both as input data. The red line denotes the mean for each method, and was used as the basis for the sorting.

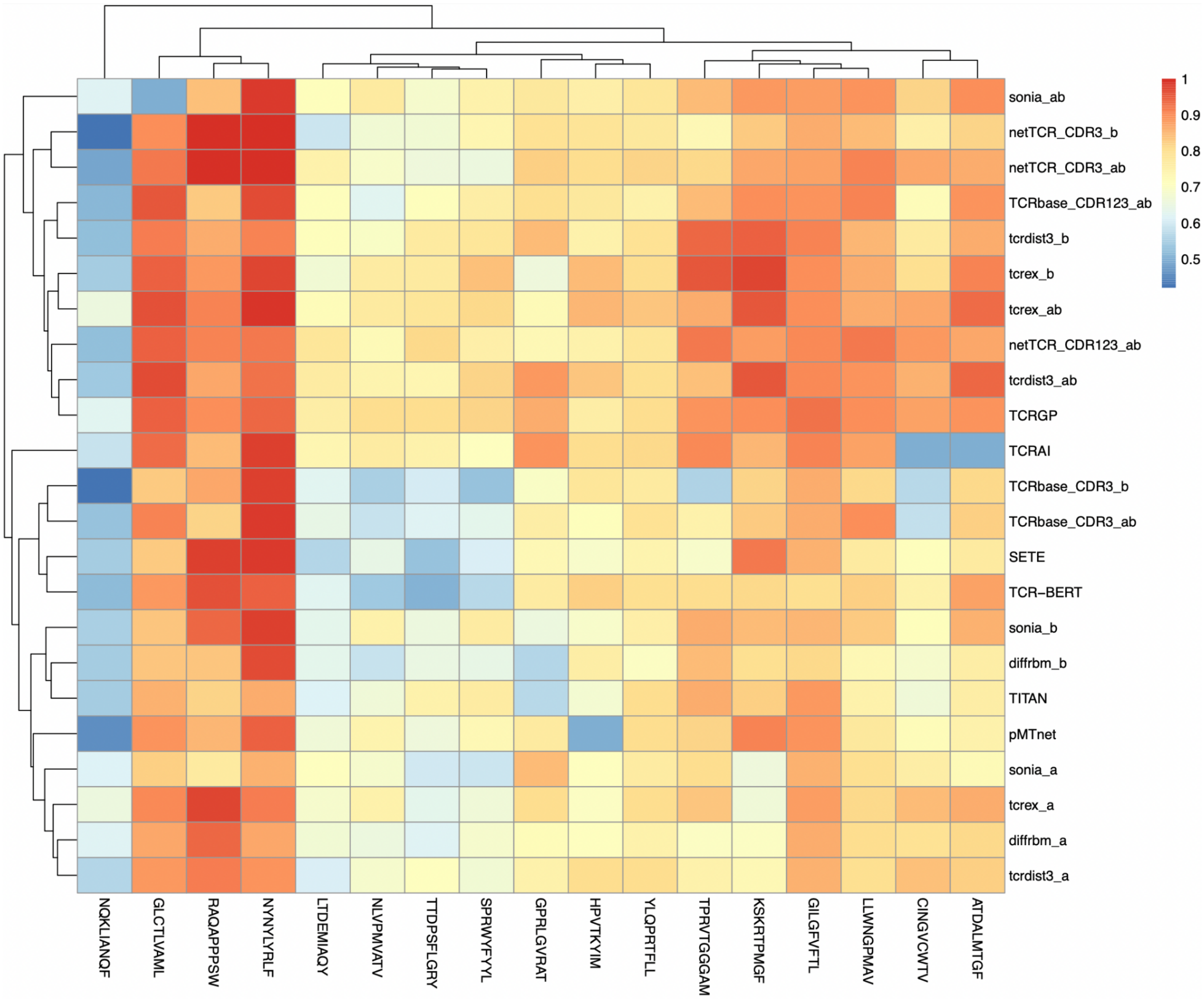


**Figure S3. Heatmap of Micro AUCs per method and epitope.** Both methods and epitopes are hierarchically clustered based on euclidean distance.

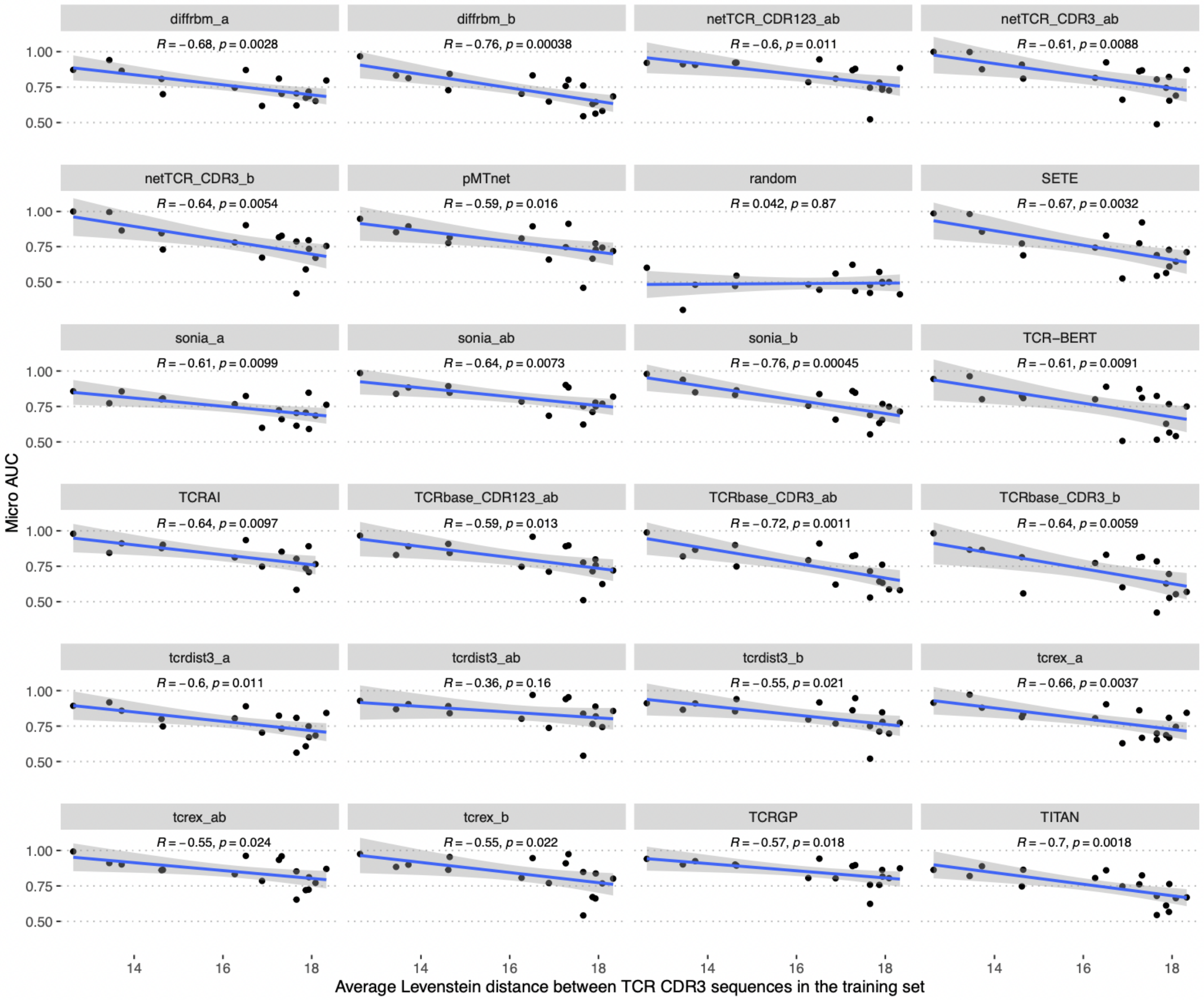


**Figure S4. Relationship between performance and the complexity of training data.** A mean pairwise distance between TCR sequences in the training sets (computed as a sum of Levenstein distances between CDR3a and CDR3b sequences) was considered as a measure of complexity of the training data. Pearson correlation coefficients of the value of this metric for different epitopes and performance of evaluated methods are shown in each plot. The only method for which p-value > 0.05 is tcrdist3_ab.
